# Dopamine and risky decision-making in pathological and problem gamblers

**DOI:** 10.1101/805929

**Authors:** Jan Peters, Taylor Vega, Dawn Weinstein, Jennifer Mitchell, Andrew Kayser

## Abstract

Gambling disorder is a behavioral addiction that is associated with impairments in value-based decision-making such as increased temporal discounting and reduced risk-aversion. Dopamine regulates learning and decision-making by modulating information processing throughout fronto-striatal circuits. Although the role of alterations in dopamine neurotransmission in the etiology of gambling disorder is controversial, preliminary evidence suggests that specifically increasing frontal dopamine levels might improve cognitive functioning in pathological and problem gamblers. We therefore examined whether increasing frontal dopamine levels via the catechol-O-methyltransferase (COMT) inhibitor tolcapone would reduce risky choice in a group of pathological and problem gamblers (n=14) in a repeated-measures counter-balanced placebo-controlled double-blind study. Choice data were fit using hierarchical Bayesian parameter estimation and a modeling scheme that combined a risky choice model with the drift diffusion model to account for both choices and response time distributions. Model comparison revealed that the data were best accounted for by a variant of the drift diffusion model with a non-linear modulation of trial-wise drift rates by value differences, confirming recent findings. Contrary to our hypothesis, risk-taking was slightly increased under tolcapone vs. placebo (*Cohen’s d* = −.281). Examination of drug effects on diffusion model parameters revealed an increase in the value-dependency of the drift rate (*Cohen’s d* = .932) with a simultaneous reduction in the maximum drift rate (*Cohen’s d* = −1.84). These results add to previous work on the potential role of COMT inhibitors in behavioral addictions, and show no consistent beneficial effect of tolcapone on risky choice in gambling disorder. Modeling results add to mounting evidence for the applicability of diffusion models in value-based decision-making. Future work should focus on individual genetic, clinical and cognitive factors that might account for the heterogeneity in the effects of COMT inhibition.

## Introduction

Gambling disorder is a prototypical behavioral addiction that constitutes a major public health concern^1^, with estimated worldwide prevalence rates ranging from 0.12 to 5.8%^2^. Both behavioral and neural similarities between gambling disorder and substance use disorders^3^ have led to the classification of gambling disorder in the category of substance-related and addictive disorders in the DSM-V^4^. Dysregulation in the dopamine system has long been implicated in substance use disorders^5,6^. Although gamblers also consistently show changes in the dopamine system^7–11^, there is considerable heterogeneity in the direction of these differences and the robustness of some of the reported effects has recently been called into question^12^.

This heterogeneity may partly explain the mixed results of past open-label and placebo-controlled trials of drugs targeting the dopamine system in patients with gambling disorder. While the combined dopamine antagonist and mood stabilizer olanzapine was not found to be superior to placebo in two studies^13,14^, both the dopamine D1 receptor antagonist ecopipam^15^ and the catechol-O-methyltransferase (COMT) inhibitor tolcapone^16^ showed promising results in open-label studies. One possible reason for these different study outcomes could be the mechanism of dopamine inactivation. In the striatum, dopamine inactivation is accomplished mainly via reuptake through the dopamine transporter. In the frontal cortex, on the other hand, this is accomplished via degradation by COMT. COMT inhibition by tolcapone is therefore thought to lead to a relatively specific increase in dopamine availability in the frontal cortex^17,18^, which may in turn augment top-down control.

Consistent with this idea, problem gambling may be more frequent in gamblers that carry the Val/Val polymorphism of the COMT val158met allele (rs4680)^19^ that codes for a more active form of the COMT enzyme, presumably leading to lower frontal dopamine levels. This association suggests that reduced frontal dopamine might increase the likelihood of problem gambling behavior. In line with such a mechanism, Grant et al. (2013) showed that the degree to which tolcapone reduced compulsivity in pathological gamblers correlated with tolcapone-induced augmentations in fronto-parietal activity^16^. Likewise, we have previously shown that tolcapone reduces impulsive choice in pathological and problem gamblers in proportion to its effect on fronto-striatal connectivity^20^. Further potential effects of tolcapone include a reduction of impulsive choice in healthy participants^21^, improved time perception^22^, increased exploration relative to exploitation^23^, and effects on working memory and risk-taking^24^.

These behavioral domains generally resonate with domains where gambling disorder is associated with impairments, in particular with respect to decision-making and executive control. A range of studies have shown increased temporal discounting (that is, a greater preference for smaller-sooner over larger-later rewards) in gamblers compared to healthy controls^25^, an effect that is also observed in substance use disorders^26^. Additionally, however, gamblers show reduced risk aversion^25^, such that they over-estimate winning probabilities during risky choice^27,28^. Risk-taking, in particular for potential gains, is increased following the administration of the dopamine precursor L-DOPA^29,30^. L-DOPA is converted into dopamine via dopamine decarboxylase, which in the human brain is expressed predominantly in the striatum^31^. L-DOPA might therefore boost dopamine availability more in the striatum than in the cortex. L-DOPA also increases temporal discounting relative to placebo^32^, whereas individual differences in dopamine availability may co-vary with temporal discounting in psychiatric populations but not healthy participants^33^.

Taken together, an increase in (presumably predominantly striatal) dopamine might increase impulsivity^32^ and risk-taking^29,30^, whereas increasing frontal dopamine levels via COMT inhibition generally tends to improve decision-making and impulse control (with potential effects of COMT genotype status^24^). Based on these observations, we examined a subset of problem and pathological gamblers from our previous randomized, double-blind, placebo-controlled crossover study^20^ to assess whether increasing frontal dopamine levels via the COMT inhibitor tolcapone would reduce risk-taking behavior. Based on recent work in reinforcement learning^34–36^, temporal discounting, and risky choice^37^ we applied a modeling framework based on the drift diffusion model^38^ in the context of a hierarchical Bayesian estimation scheme. This modeling approach has the benefit of accounting for the full response time (RT) distributions associated with decisions, thereby providing more detailed information regarding choice dynamics^34^.

## Methods

### Participants

Participants were recruited via online advertisements (Craigslist) and screened. Subjects with South Oaks Gambling Screen (SOGS)^39^ scores > 5 were considered eligible and invited to participate. Table 1 provides an overview of the clinical and demographic data of all participants. The study procedure was approved by the local institutional review board and participants provided written informed consent prior to participation.

**Table 1.** Demographic and clinical characteristics of the problem and pathological gamblers. ^a^South Oaks Gambling Screen^39^,^b^Gambling-Related Cognitions Scale^40^, ^c^Beck Depression Inventory^41^, ^d^Alcohol Use Disorders Identification Test^42^, ^e^Barratt Impulsivity Scale^43^.

|  | N / Mean (SD) | Range |
| --- | --- | --- |
| **N female/male** | 6/8 |  |
| **N Smokers/Nonsmokers** | 4/10 |  |
| **COMT genotype**<br>**(Val/Val, Val/Met, Met/Met)** | 7/4/3 |  |
| **Age** | 32.57 (9.03) | 20-47 |
| **YoE** | 14.93 (1.86) | 12-18 |
| **SOGS<sup>a</sup>** | 10.79 (3.07) | 6-18 |
| **GRCS<sub>Total</sub><sup>b</sup>** | 97.79 (14.08) | 76-116 |
| **BDI<sup>c</sup>** | 11.79 (7.89) | 0-27 |
| **AUDIT<sup>d</sup>** | 11.93 (6.40) | 2-20 |
| **BIS<sup>e</sup>** | 70.5 (9.62) | 50-88 |

As described in our previous study^20^, subjects were required to be between 18 and 50 years old, in good health, able to read and speak English, and able to provide informed consent. Women of reproductive age were required to be using an effective form of contraception, and to be neither pregnant nor lactating during study participation. A positive urine drug toxicology screen before any visit was grounds for exclusion, as was an alcohol level greater than zero as measured by breathalyzer before any visit. Similarly, subjects were excluded for reported use of psychoactive substances (including both prescription medications and drugs of abuse) within the prior two weeks, use of drugs of abuse more than ten times in the previous year, or current dependence on marijuana. Subjects could otherwise use marijuana no more than three times per week and were required to refrain from marijuana use for at least 48 hours prior to testing sessions. These criteria did not apply to nicotine; the two regular smokers (out of four total nicotine-using subjects) were both easily able to refrain for the duration of specific study sessions. Subjects who were taking medications with dopaminergic, serotonergic, or noradrenergic actions (although animal work suggests that tolcapone induces increases in dopaminergic but not noradrenergic concentrations^18^) or who had a known allergy to either tolcapone or the inert constituents in tolcapone capsules, were also excluded. Likewise, after completion of the Mini International Neuropsychiatric Interview^44^, subjects who met screening criteria for an axis I psychiatric disorder other than gambling disorder, such as major depression, or who had a significant medical or psychiatric illness requiring treatment, were excluded from participating. Because tolcapone carries the potential for hepatotoxicity, liver function tests as assessed by phlebotomy were required to be no more than three times the upper limit of normal.

### Drug administration

Subjects were randomized in double-blind, counterbalanced, placebo-controlled fashion to either placebo or a single 200mg dose of tolcapone on their first visit and the alternative treatment on their second visit. This dose was based upon our previously published findings that a single 200mg dose has measurable behavioral effects^21,23,45^. Subjects began the current task approximately 3 hours after tolcapone and placebo ingestion, following the MRI procedures described in our previous study^20^. Tolcapone is expected to have pharmacodynamically relevant serum concentrations for at least 6 hours^46,47^. No subjects reported potential side effects under either the placebo or tolcapone conditions during their participation.

### Risk-taking task

On each testing day, participants completed 96 trials of a risky-choice task involving a series of choices between a smaller, certain reward ($10 with 100%) and larger, but riskier, options. A first set of risky options consisted of all combinations of sixteen reward amounts (10.1, 10.2, 10.5, 11, 12, 15, 18, 20, 25, 30, 40, 50, 70, 100, 130, 150 dollars) and seven probabilities (10%, 17%, 28%, 54%, 84%, 96%, 99%). We used a second set of probabilities (11%, 18%, 27%, 55%, 83%, 97%, 98%) in combination with the same series of reward amounts to create a second set of 96 trials. The assignment of the two sets of trials to the two drug conditions was randomized across participants. The experiment was implemented in Presentation^©^ (Neurobehavioral Systems). Trials were presented in randomized order and with a randomized assignment of safe/risky options to the left/right side of the screen. Both options remained on the screen until a response was made.

### Computational modeling

#### Risky choice model

We applied a simple single-parameter discounting model to describe how value changes as a function of probability, such that discounting is hyperbolic over the odds against winning the gamble^48^:

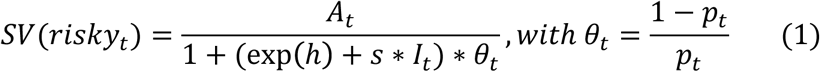

Here, *A* is the numerical reward amount of the risky option, θ is the odds against winning and *I* is an indicator variable that takes on a value of 1 for tolcapone data and 0 for placebo data. The model has two free parameters: *h* is the hyperbolic discounting rate from the placebo condition (modeled in log-space) and *s* is a weighting parameter that models the degree of reduction in discounting under tolcapone vs. placebo.

### Softmax action selection

Softmax action selection models the choice probabilities as a sigmoid function of value differences^49^:

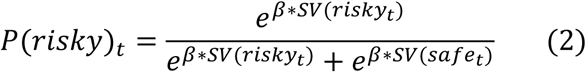

Here, *SV* is the subjective value of the risky reward according to Eq. 1 and *β* is an inverse temperature parameter, modeling choice stochasticity (for *β* = 0, choices are random and as *β* increases, choices become more dependent on the option values).

### Drift diffusion choice rule

To better characterize the dynamics of the decision process, we replaced the softmax choice rule (Eq. 2) with the drift diffusion model (DDM), based on recent work in reinforcement learning^34–36^. The DDM accounts not only for binary choices but for the full reaction time distributions associated with those decisions. We used the Wiener Module^50^ for the JAGS statistical modeling package^51^ that implements the likelihood function of a Wiener diffusion process. The DDM assumes that decisions arise from a noisy evidence accumulation process that terminates as the accumulated evidence exceeds one of (usually) two decision bounds. Reinforcement learning applications of the DDM have used accuracy coding to define the response boundaries of the DDM^34–36^, such that the upper boundary corresponds to selections of the objectively superior stimulus, and the lower boundary to choices of the inferior option. This structure is in line with the traditional application of the DDM in the context of perceptual decision-making tasks^52^. However, in value-based decision-making, there is typically no objectively correct response. Therefore, previous applications of the DDM in this domain have instead re-coded accuracy to correspond to the degree to which decisions are consistent with previously obtained preference judgements^53^. This approach is not possible, however, when the goal is to use the DDM to model the preferences that in such a coding scheme would determine the boundary definitions. Therefore, here we applied stimulus coding, such that the upper boundary (1) corresponded to the selection of the risky option and the lower boundary (0) to the selection of the certain option.

We used percentile-based cut-offs for RTs, such that for each participant, the fastest and slowest 2.5% of trials were excluded. RTs for choices of the certain 100% option were then multiplied by −1 prior to model estimation. The RT on a given trial is then distributed according to the Wiener First Passage Time (WFPT):

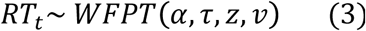

Here, *α* is the boundary separation (modeling response caution / the speed-accuracy trade-off), *z* is the starting point of the diffusion process (modeling a bias towards one of the decision boundaries), *τ* is the non-decision time (reflecting perceptual and/or response preparation processes unrelated to the evidence accumulation process) and *v* is the drift rate (reflecting the rate of evidence accumulation). In the JAGS implementation of the Wiener model^50^, the starting point *z* is coded in relative terms and takes on values between 0 and 1. That is, *z* = .5 reflects no bias, *z* >.5 reflects a bias towards the upper (risky option) boundary, and *z* <.5 reflects a bias towards the lower (certain option) boundary.

We then compared three variants of the DDM: First, we examined a null model (DDM_0_) without any value modulation. In this model, the four DDM parameters (*α*, *τ*, *z*, and *v*) were held constant across trials. Drug effects were modeled by including a term modeling a tolcapone-induced change relative to the placebo condition for each parameter. Second, we examined two previously proposed functions linking trial-by-trial changes in the drift rate *v* to value-differences. We examined a linear mapping (DDM_lin_) as proposed by Pedersen et al. (2017)^34^:

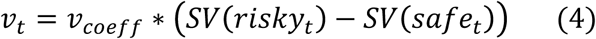

Here, *v*_*coeff*_ maps trial-wise value differences onto the drift rate *v*. *SV* is the subjective value of the rewards according to Eq. 1.

We also examined a recently proposed non-linear (DDM_S_) scheme^35^:

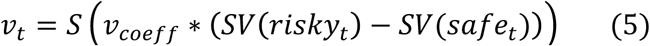

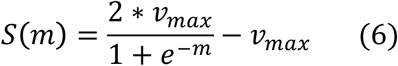

Here, *S* is a sigmoid function centered at 0 with *m* being the scaled value difference from Eq. 5, and asymptote ± *v*_*max*_. For DDM_lin_ and DDM_S_, effects of choice difficulty on response times naturally arise. For more similar values, the trial-wise drift rate approaches 0.

### Hierarchical Bayesian models

Model building proceeded as follows. As a first step, all models were fit at the level of individual participants. We validated that good fits could be obtained, such that posterior distributions were centered at sensible parameter values and the Gelman-Rubin 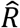 statistic, an estimate of the degree of Markov chain convergence (see below), was in an acceptable range of 1 ≤ 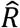 ≤ 1.01. In a second step, models were fit in a hierarchical manner with group-level distributions for all parameters. We used the same convergence criteria as for the single-subject models (1 ≤ 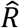 ≤ 1.01). For group level hyper-parameters, we used uninformative priors (i.e. uniform distributions for means defined over sensible ranges, gamma distributions for precision). Here, models were fit separately to the data from the placebo and tolcapone conditions, to examine whether drug administration altered the relative model ranking. Finally, after identifying the variant of the drift diffusion model that accounted for both the placebo and tolcapone data best, we fit this model across drug conditions. In this final combined model, parameters from the placebo condition were modeled as the “baseline”, and all drug effects were modeled as Gaussians with group level priors with *μ* = 0, *σ* = 2. All JAGS model code is available on the Open Science Framework (https://osf.io/wtg89/).

### Model estimation and comparison

Models were fit using Markov Chain Monte Carlo (MCMC) as implemented in JAGS^51^ (Version 4.2) with the *matjags* interface (https://github.com/msteyvers/matjags) for Matlab^©^ (Mathworks) and the JAGS Wiener module^50^. For each model, we ran two chains with a burn-in period of 100k samples and thinning of 2. 10k additional samples were then retained for further analysis. Chain convergence was assessed via the 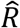 statistic, where we considered 1 ≤ 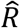 ≤ 1.01 as acceptable values for all group- and individual-level parameters. Relative model comparison was performed via the Deviance Information Criterion (DIC), where lower values indicate a better fit^54^.

### Posterior predictive checks

We additionally performed posterior predictive checks to ensure that the best-fitting model captured key aspects of the data. Therefore, during model estimation, we simulated 10k full datasets from the hierarchical models based on the posterior distribution of parameters. For each participant and drug condition, model-predicted RT distributions for a random sample of 1k of these simulated data sets were then smoothed with non-parametric density estimation (*ksdensity.m* in Matlab) and overlaid on the observed RT distributions for each subject and drug condition.

### Analysis of drug effects

We characterize drug effects in the following ways. First, we show group posterior distributions for all parameters, and 85% and 95% highest density intervals for the posterior distributions of the tolcapone-induced changes in parameters (shift parameters). Additionally, we report Bayes Factors for directional effects^34,55^ based on the posterior distributions of these shift parameters. This value was determined via non-parametric kernel density estimation in Matlab (*ksdensity.m*) and computed as *BF* = *i*/(1 − *i*), where *i* is the integral of the posterior distribution from 0 to +∞. Following common criteria, Bayes Factors >3 indicate support for a model, whereas Bayes Factors >12 indicate substantial support. Conversely, Bayes Factors <.33 are interpreted as evidence in favor of the alternative model. Lastly, we report standardized effect sizes for all drug-induced changes, which we calculated based on the means of the group-level posterior mean and precision parameters of the hierarchical model.

### Genetics

DNA extraction and SNP analysis were performed by the Genomics Core at the UCSF Institute for Human Genetics on salivary samples (salimetrics.com) collected during the screening visit. DNA was extracted using Gentra Puregene reagents and protocols and quantified using the Pico Green method (Molecular Probes/Invitrogen). Genotyping of the cathechol-*O*-methyltransferase (COMT; rs4680) polymorphism via polymerase chain reaction was carried out using TaqMan^®^ technology (Applied Biosystems).

## Results

### Model-free analyses

Response time (RT) distributions across participants per drug condition are shown in Figure 1a. Arcsine-square-root transformed risky choice ratios (Figure 1b) did not differ significantly between drug conditions (t_13_=-.677, p=.51, 95% CI: [-.18, .095]). Likewise, median RTs did not differ significantly between drug conditions (t_13_=-.184, p=.857, 95% CI: [-.32, .27]).

**Figure 1.**
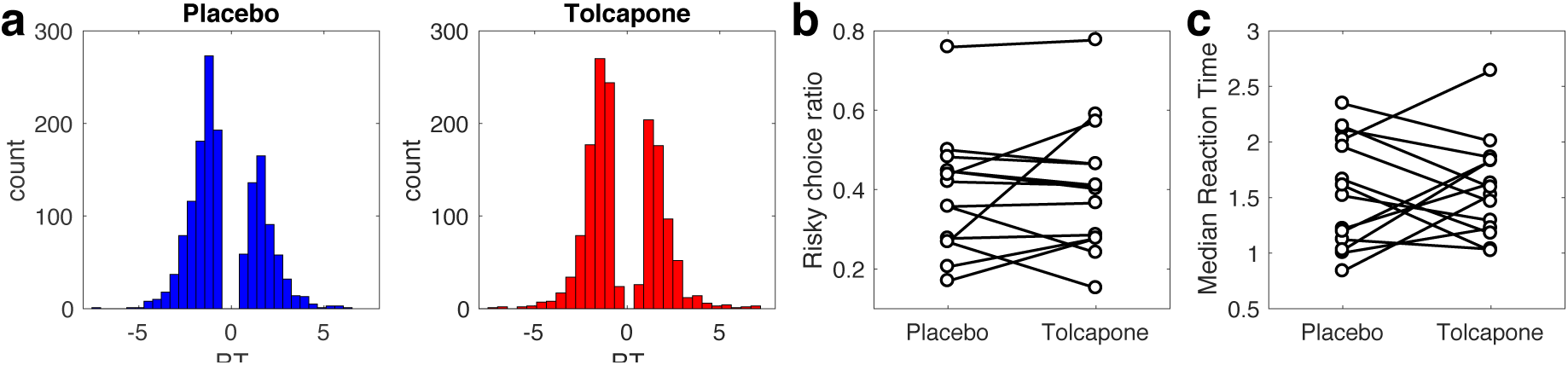
a) Overall RT distributions for placebo (blue) and tolcapone (red). b) Proportion of choices of the risky option per participant and drug-condition. c) Median RT per participant and drug condition.

### Model comparison

We then compared three variants of the DDM: a null model without any value modulation (DDM_0_), a model with a linear scaling of trial-wise drift rates (DDM_lin_) and a model with non-linear (sigmoid) drift rate scaling (DDM_S_). To ensure that drug condition did not impact model ranking, we first fit the three models separately to the data from the placebo and tolcapone conditions. As can be seen from Table 2, model ranking was the same in the two drug conditions, such that models including value modulation of the drift rate outperformed the DDM_0_, and the non-linear DDM_S_ fit the data better than the DDM_lin_.

**Table 2.** Model comparison of the drift diffusion models, separately for the two drug conditons. Under both placebo and tolcapone, the data were best accounted for by a model including a non-linear mapping from trial-wise value-differences to drift rates (DDM_S_).

| Model | Placebo |  | Tolcapone |  |
| --- | --- | --- | --- | --- |
|  | DIC | Rank | DIC | Rank |
| $DDM_0$ | 42383 | 3 | 43177 | 3 |
| $DDM_{lin}$ | 36136 | 2 | 38302 | 2 |
| $DDM_s$ | 30354 | 1 | 32240 | 1 |

### Initial model validation

Initial analyses confirmed that risk-taking behavior quantified via softmax action selection could reliably reproduced when using the DDM_S_ as the choice rule (see Supplemental Figure 1).

### Posterior predictive checks

We next fit the DDM_S_ to the combined data from the two drug conditions, modeling the placebo condition as the baseline, and tolcapone-induced changes in each parameter as additive changes relative to that baseline (see methods) using Gaussian priors centered at zero. Then we examined the extent to which the DDM_S_ could reproduce the reaction time distributions observed in individual participants. To this end, we simulated 10k full datasets from the models’ posterior distribution. The histograms in Figure 1 show the observed reaction time distribution for each participant and drug condition, with a smoothed density estimate of the model-generated reaction time distribution (based on 1000 random samples from the simulations) overlaid. Generally, the model accounted reasonably well for the observed reaction time distributions in most participants. The DDM_S_ also accounted for a similar proportion of binary decision under tolcapone and placebo (M[range]_placebo_: .899 (.798-.962), M[range]_tolcapone_: .879 (.717-.972), t_13_=1.21, p=.249).

**Figure 2.**
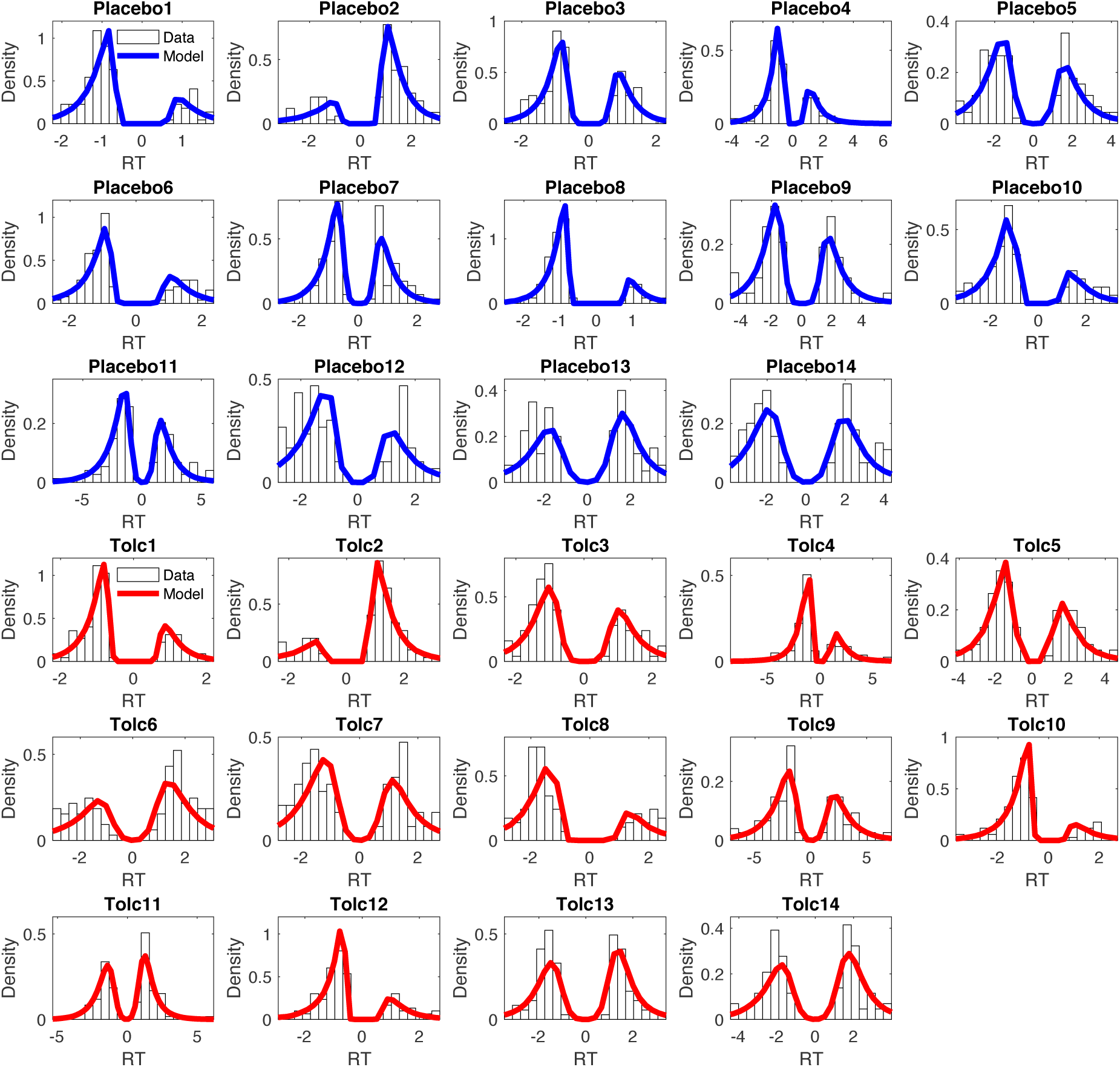
Posterior predictive plots of the drift diffusion temporal discounting model with non-linear value scaling of the drift rate (DDM_S_) for all fourteen participants (blue: placebo, red: tolcapone). Histograms depict the observed RT distributions for each participant. The solid lines are smoothed histograms of the model predicted RT distributions from 1k individual subject data sets simulated from the posterior of the best fitting hierarchical model. RTs for smaller-sooner choices are plotted as negative, whereas RTs for larger-later choices are plotted as positive. The x-axes are adjusted to cover the range of observed RTs for each participant.

**Table 2.**
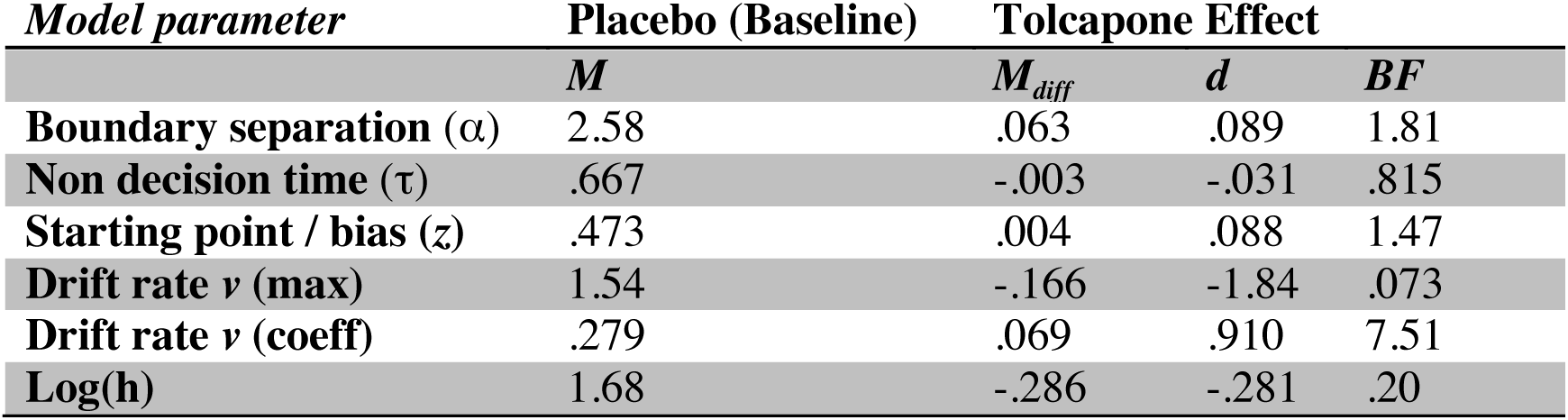
Summary of drug effects on model parameters: For each parameter, we report the mean estimate from the placebo (baseline) condition, the mean parameter changes under tolcapone, standardized effect sizes (Cohen’s *d*) that were calculated based on the posterior group-level estimates of mean and precision (see methods section) and Bayes Factors (BF) testing for directional effects^34,55^. Bayes Factors <.33 indicate evidence for a reduction under tolcapone, whereas Bayes Factors >3 indicate evidence for an increase under tolcapone (see Methods section).

| *Model parameter* | **Placebo (Baseline)** | **Tolcapone Effect** |  |  |
| --- | --- | --- | --- | --- |
| | $M$ | $M_{diff}$ | $d$ | $BF$ |
| **Boundary separation ($\alpha$)** | 2.58 | .063 | .089 | 1.81 |
| **Non decision time ($\tau$)** | .667 | -.003 | -.031 | .815 |
| **Starting point / bias ($z$)** | .473 | .004 | .088 | 1.47 |
| **Drift rate $v$ (max)** | 1.54 | -.166 | -1.84 | .073 |
| **Drift rate $v$ (coeff)** | .279 | .069 | .910 | 7.51 |
| **Log(h)** | 1.68 | -.286 | -.281 | .20 |

### Effects of tolcapone on risk-taking and diffusion model parameters

We next examined the posterior distributions of parameters of the final DDM_S_ model in more detail. Figure 3 (top row) shows the group level posterior distributions for parameters at baseline (placebo) as well as for the tolcapone effects (Figure 3, bottom row). For comparison only, Figure 3 additionally shows posterior distributions from our previous study^37^ using the same task in medial orbitofrontal cortex lesion patients (n=9) and controls matched on age- and education to the lesion patients (n=19). Under placebo, both *response caution* (boundary separation) and *non-decision time* in the gamblers were substantially lower than the corresponding values in these two groups. Gamblers also exhibited a bias towards the safe option, reflected in a posterior distribution of the starting point that was shifted slightly towards zero (Fig. 3c), but was numerically more similar to the mOFC patient group than to the control group. The maximum drift rate *v*_*max*_ at placebo was numerically higher in the gamblers (Fig. 3d) and there was a robust positive effect of value-differences on the trial-wise drift rates, as reflected in a positive drift rate coefficient parameter under placebo (*v*_*coeff*_, Fig. 3e). Interestingly, *log(h)* (i.e. risk-taking) in the gamblers under placebo (Fig. 3f) was qualitatively more similar to risk-taking behavior of the mOFC patients than that of the controls from our previous study.

**Figure 3.**
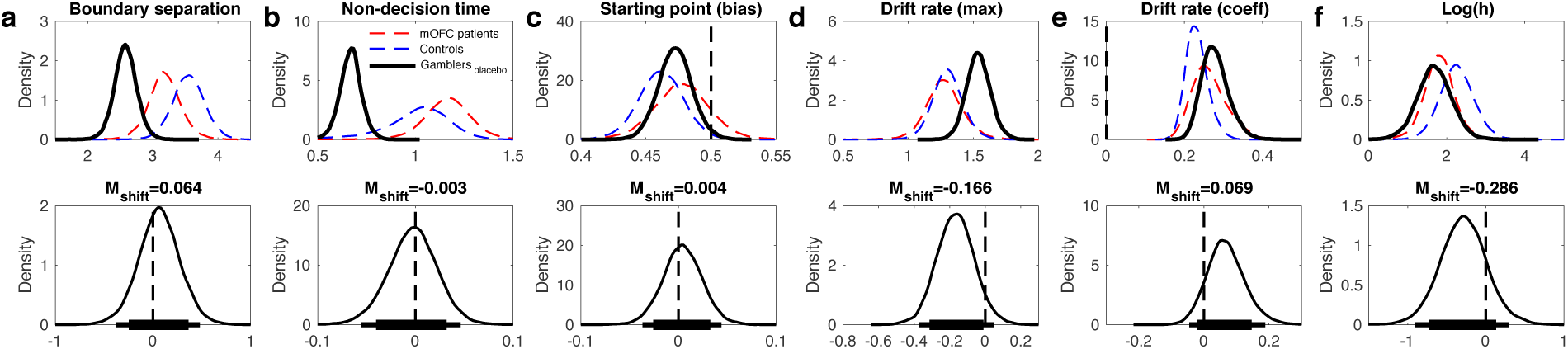
Top row: Group-level posterior distributions for parameter means under placebo (solid black line, a: boundary separation, b: non-decision time, c: bias, d: vmax, e: vcoeff, f: log(h) [risk-taking]). For comparison but not inference, the dashed lines plot the group posterior distributions from our previous study using the same task^37^ (red: n=9 mOFC lesion patients, blue: n=19 matched controls). Bottom row: group level posterior distributions for tolcapone-induced changes for each parameter. The thin solid lines in the bottom row indicate 95% highest density intervals, whereas the thick solid horizontal lines indicate 85% highest density intervals.

All drug effects are summarized in Table 2 (mean parameter changes between tolcapone and placebo, standardized effect sizes (Cohen’s *d*), Bayes Factors for directional effects; see methods section). The posterior distributions for the tolcapone-induced change for boundary separation (Fig. 3a), non-decision time (Fig. 3b) and starting point (Fig. 3c) were all centered at zero with effect sizes of |*d*|<.1. In contrast, under tolcapone, there was evidence for a decrease in the maximum drift rate (*v*_*max*_) (*d* = −1.84, BF = .073), an increase in the value-dependent drift-rate modulation (*d* = .901, BF = 7.51) and for an increase in risk-taking (*d* = −.281, BF = .20).

### Consistency of tolcapone effects across participants

We next examined the consistency of the latter three group effects across participants by overlaying individual posterior distributions for the tolcapone-effects over the average group effects for parameters showing drug-effects at the group level (Figure 4: *v*_*max*_, Figure 5: *v*_*coeff*_, Figure 6: *log(h)*). Under tolcapone, 13/14 participants showed and a mean reduction in the maximum drift rate *v*_*max*_, 12/14 showed an increase in the drift rate scaling *v*_*coeff*_, and 9/14 showed an decrease in *log(h)* (increase in risk-taking). For transparency, we have highlighted the three Met/Met genotype participants in these plots (red lines), being fully aware that interpretation of genotype effects in such a small sample is unwarranted

**Figure 4.**
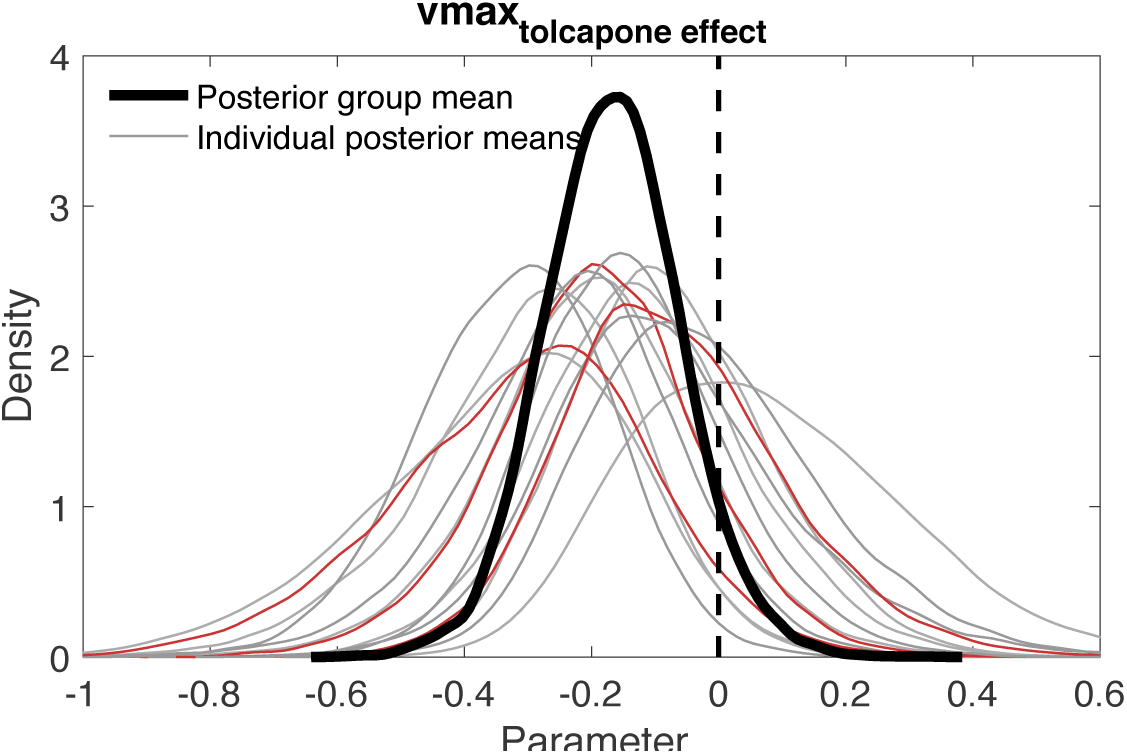
Posterior group mean (solid black line) for the tolcapone-induced change in maximum drift rate (*v*_*max*_, see Eq. 6) and individual subject posterior distributions of the same effect (grey: Val/Val and Val/Met, red: Met/Met). The mean change was < 0 in 13/14 subjects.

**Figure 5.**
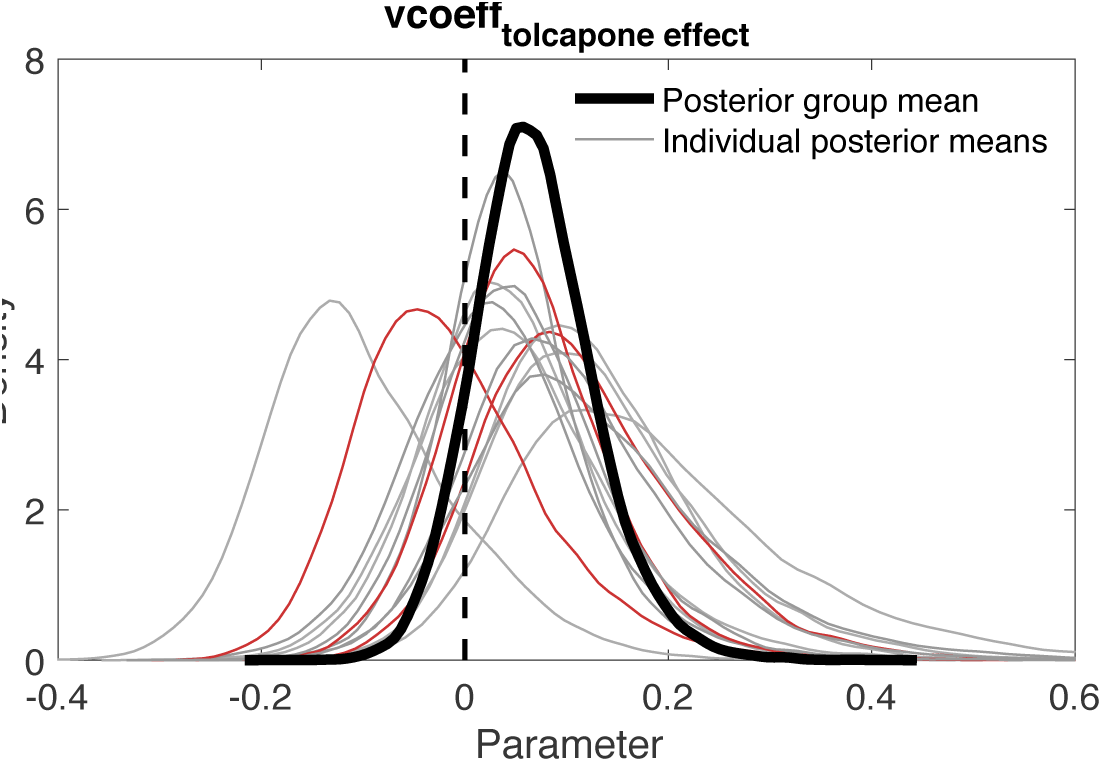
Posterior group mean (solid black line) for the tolcapone-induced change in value-dependent drift rate modulation (v_coeff_, see Eq. 5) and individual subject posterior distributions of the same effect (grey: Val/Val and Val/Met, red: Met/Met). The mean change was > 0 in 12/14 subjects.

**Figure 6.**
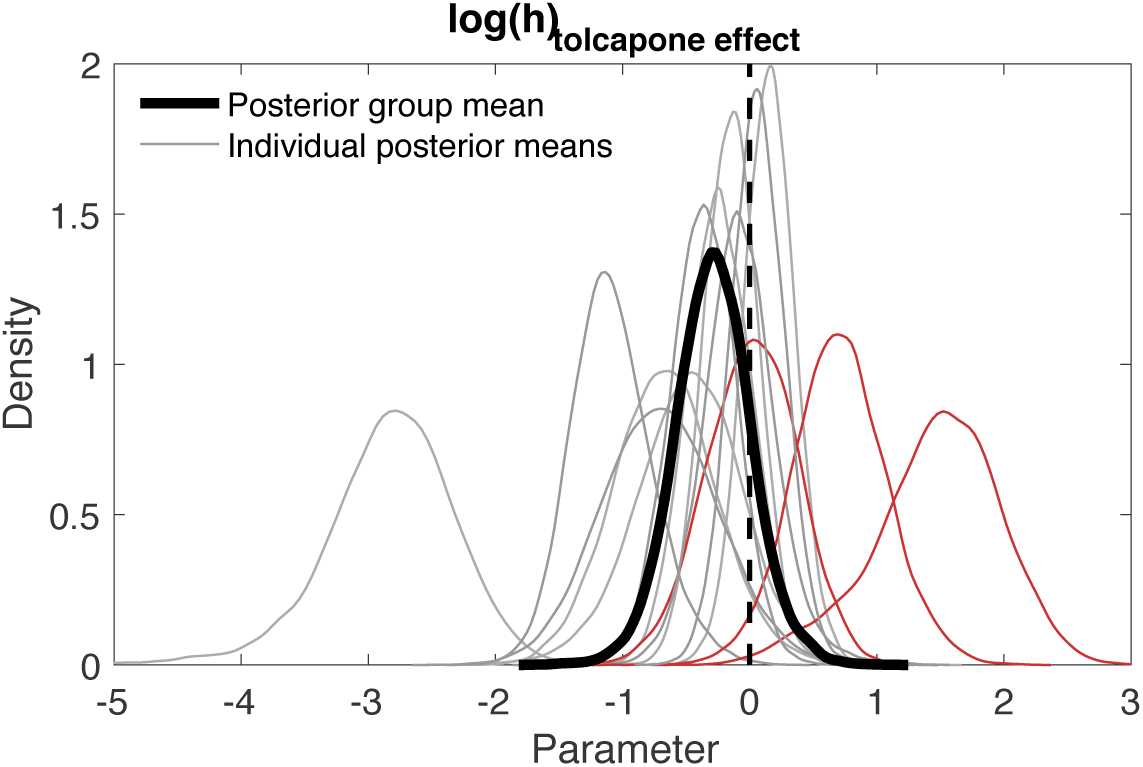
Posterior group mean (solid black line) for the tolcapone-induced change in risk-taking (log(h), see Eq. 1) and individual subject posterior distributions of the same effect (grey: Val/Val and Val/Met, red: Met/Met). The mean change was < 0 in 9/14 subjects.

## Discussion

Gambling disorder is associated with impairments in value-based decision-making, including increased temporal discounting and reduced risk aversion^25^. Here we tested whether risk-taking in problem and pathological gamblers could be attenuated by the COMT inhibitor tolcapone, which predominantly increases dopamine levels in the frontal cortex. Choice data were modeled in a hierarchical Bayesian scheme with the drift diffusion model as the choice rule to account for both choices and reaction time distributions. In contrast to our initial hypothesis, if anything tolcapone increased risk-taking (small effect size). Examination of the drift diffusion model parameters showed a reduction in the maximum drift rate under tolcapone (large effect size) and an increase in the value-dependency of the drift rate (large effect size).

We used a modeling scheme based on the drift diffusion model, which has recently gained some popularity in reinforcement learning and value-based decision-making^34–37^. As in our previous work^37^, choice model parameters estimated via a standard softmax function could be reliably reproduced using the diffusion model as the choice rule. Posterior predictive checks revealed that the best fitting drift diffusion model reproduced individual subject reaction time distributions reasonably well in both drug conditions. In keeping with previous work^35,37^, we again carried out a model comparison and evaluated both a linear and non-linear mapping from value-differences to trial-wise drift rates. The DDM_S_ fit the data better in both drug conditions, confirming previous results in non-linear drift rate scaling.

Our results suggest small effects (|*d*|<.1) of tolcapone on three parameters of the drift diffusion model: boundary separation, non-decision time and starting point (bias). This finding suggests that overall response caution (as reflected in the boundary separation parameter) and processes related to motor preparation and/or stimulus processing (as reflected in the non-decision time) were largely unaffected by tolcapone. Since group differences for these parameters are potentially confounded by age, we do not discuss them further. Likewise, gamblers under placebo showed a bias towards the safe option (*z*_*placebo*_ = .473, see Figure 3c) that was similar to the bias exhibited by both medial orbitofrontal cortex lesion patients and controls in our previous study using the same risk-taking task^37^ (*z*_*mOFC*_ = .478, *z*_*controls*_ = .461), but numerically closer to the patient’s bias. Note that gamblers under placebo also exhibited a level of risk-taking that was numerically even more pronounced than that of the medial orbitofrontal cortex lesion patients from our previous study^37^ (*log(h)*_placebo_ = 1.68, *log(h)*_mOFC_ = 1.81, *log(h)*_controls_ = 2.26). Under tolcapone, this risk-taking increased even further (*log(h)*_tolcapone_ = 1.39). Despite the absence of a specific control group for the present sample of pathological and problem gamblers, this finding nonetheless suggests that risk-taking behavior was quite pronounced in the gamblers, and puts the drug-effect in perspective. Taken together with the observation regarding the bias, this further indicates that risk-taking might be generally associated with reduced bias for the safe option.

What mechanism might drive the observed effects of tolcapone on risk-taking and value evidence accumulation? Our approach was motivated by the idea that tolcapone might attenuate risk-taking via an augmentation of prefrontal cortex (top-down control) functions. The lateral prefrontal cortex is implicated in cognitive control^56,57^, and disruption of prefrontal cortex function can increase risk-taking and impulsivity^37,58–61^. Likewise, tolcapone has been shown to act through an enhancement of prefrontal cortex activation and /or fronto-striatal interactions^16,20,21^. However, although the drug-effect on risk-taking was small, it was in the opposite direction, increasing risk-taking rather than attenuating it. Furthermore, the directionality and effect size of the drug-effect on risk-taking showed some heterogeneity across participants (Figure 6). In the absence of task-related imaging data, drawing definite conclusions regarding the mechanism underlying these differential effects of tolcapone on risk-taking remains speculative, and individual genetic differences likely contribute to these variable results.

Similarly, it remains unclear through what exact mechanism an increase of frontal dopamine levels might affect the changes in value-dependency of the drift-rate observed in the present study. Ventromedial prefrontal cortex is involved in coding for reward valuation during learning and decision-making^62,63^. It could thus be speculated that tolcapone might enhance such value representations, thereby increasing the value-dependency of trial-wise drift rates. However, at the same time maximum drift rates were reduced under tolcapone, an effect that was consistent across participants (see Figure 4). This might reflect at a trade-off between vmax and vcoeff parameters in the model, such that reduced vmax can be compensated for by increases in vcoeff in some conditions. Such interactions require further study in the use of diffusion model based choice rules.

Finally, dopamine has different functions in different prefrontal cortex subregions^64^, such that different dopamine-dependent cognitive functions might exhibit different dose-response functions^65^ and thus be differentially modulated by tolcapone. A thorough assessment of these complexities, including process-dependent baseline effects and potential subregion-specific effects of tolcapone will need to be more fully addressed in future studies.

While we genotyped participants for the COMT Val158Met polymorphism, drawing any conclusions regarding genotype effects in a small sample study such as the present one is obviously highly problematic. On the other hand, not reporting genotype data that is available would also seem inappropriate given the previously suggested COMT genotype-dependency of tolcapone effects on risk-taking^24^. In their between-subjects study, Farrell and colleagues^24^ reported increased risk-aversion in Val/Val participants under tolcapone, compared to a group of Met/Met carriers. In contrast to that study, in our data set the two participants showing the largerst reduction in risk-taking under tolcapone were Met/Met carriers. This result is in line with the frequent observation that dopamine effects on cognitive functions mediated by the prefrontal cortex depend on baseline dopamine availability in an inverted-U-shaped fashion^66^. Yet, in this model, Met/Met carriers exhibit a higher frontal dopamine level at baseline due to the COMT enzyme being less active. Further COMT suppression (e.g. via tolcapone) is then thought to move Met/Met subjects into an “overdosed” state, impairing performance relative to placebo^24,66,67^. This is not compatible with the substantial reduction in risk-taking observed for 2/3 Met/Met carriers. However, as mentioned above, different functions might show different functional forms of dopamine baseline-dependency^62^, which would require much larger subject numbers to fully evaluate.

There are several additional limitations of the present study that need to be acknowledged. First, given the small sample size, our findings require replication in larger samples and groups other than pathological and problem gamblers. Second, although gender was relatively balanced in the present study, which is often not the case in studies involving pathological and problem gamblers, we were obviously underpowered to examine sex differences. Third, we did not test a control group specifically matched to the gamblers. Rather, we focused exclusively on potential drug-effects in a group of pathological and problem gamblers. The aim of the project was to examine the degree to which behavioral markers of gambling disorder such as risk-taking and temporal discounting^20^ could be improved by COMT inhibition, but future studies could benefit from a more detailed exploration of the effects of COMT inhibition on risk-taking in healthy controls, as previously done for inter-temporal choice^21^. However, to provide some reference for the level of risk-taking behavior in our particular sample of pathological and problem gamblers, we have plotted parameters for a group of medial orbitofrontal cortex lesion patients and controls from a previous study^37^.

Taken together, our data extend previous investigations of modeling schemes that build on the drift diffusion model^34–37^, by successfully applying this approach for the first time in a group of problem and pathological gamblers. While the data are preliminary given the small sample size, they suggest that tolcapone might impact aspects of value evidence accumulation during risky choice. However, our data do not support the idea that tolcapone consistently attenuates risk-taking in pathological and problem gamblers. These results extend and complement previous examinations of the potential of COMT inhibition in gambling disorder^16,20^ by providing a comprehensive model-based analysis of risky decision-making.

## Acknowledgements

This work was supported by the National Center for Responsible Gaming (A.S.K.) and Deutsche Forschungsgemeinschaft (DFG, Grant PE1627/5-1 to J.P.).

## Supplemental material

### Comparison with standard softmax action selection

As in our previous report^37^, we also checked whether the choice model parameters estimated via a standard softmax choice rule (see Methods section, Equation 2) could be reproduced using the DDM. We therefore correlated single subject mean posteriors for *log(h)* (risk taking under placebo) and *log(h)*_*tolceffect*_ (the change in risk taking under tolcapone) from the hierarchical DDM_S_ and the hierarchical model with softmax action selection (see Figure 2). Both parameters were highly correlated between estimation schemes (*log(h)*: *r*=.98, *p*<.0001, log(h)_tolceffect_: *r*=.93, *p*<.0001), indicating that parameters estimated via standard methods could be reproduced using the DDM^37^.

**Supplemental Figure 1.**
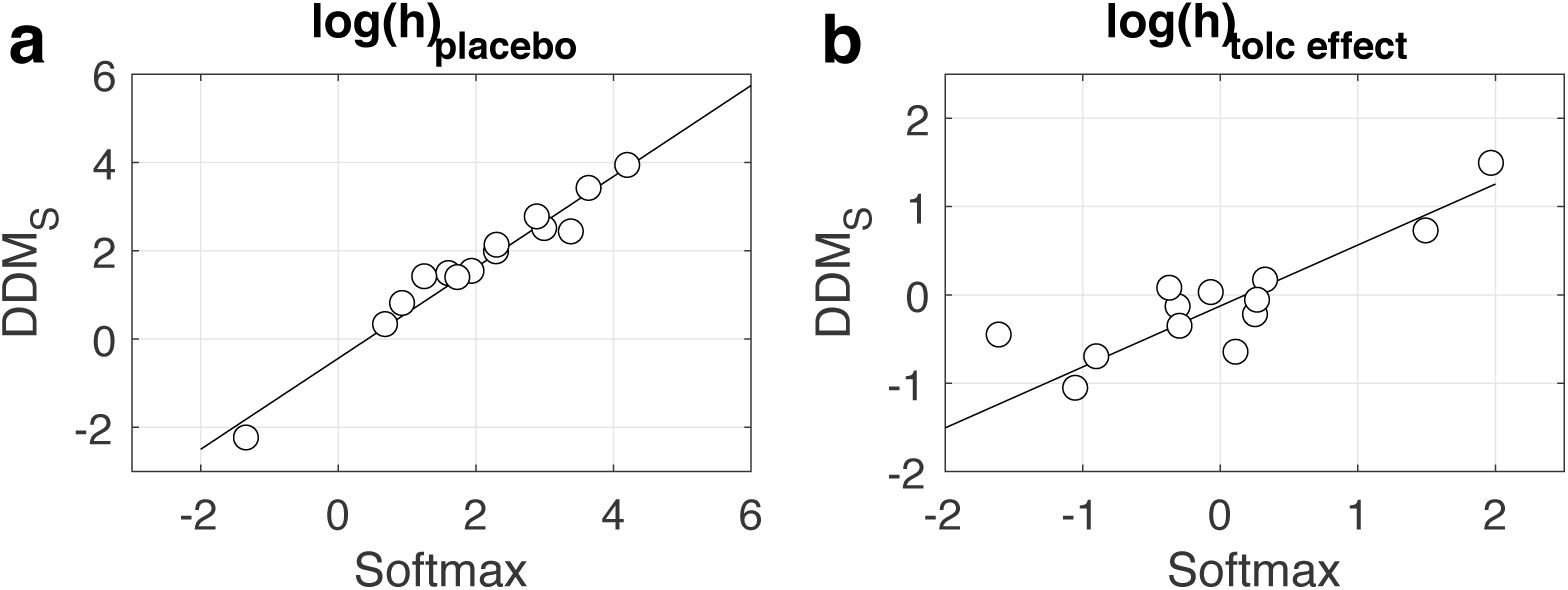
a) Correlation between the probability discount rate *log(h)* under placebo, estimated via standard softmax and via the DDM_S_. b) Correlation between the change in *log(h)* under tolcapone, estimated via standard softmax and via the DDM_S_.

